# A unique arginine cluster in PolDIP2 enhances nucleotide binding and DNA synthesis by PrimPol

**DOI:** 10.1101/2020.05.18.101550

**Authors:** Kazutoshi Kasho, Gorazd Stojkovič, Cristina Velázquez-Ruiz, Maria Isabel Martínez-Jiménez, Timothée Laurent, Aldo E. Pérez-Rivera, Louise Jenninger, Luis Blanco, Sjoerd Wanrooij

**Author notes:** To whom correspondence should be addressed: Sjoerd Wanrooij, Department of Medical Biochemistry and Biophysics, Umeå University, 90187 Umeå, Sweden.

## Abstract

Replication forks often stall at damaged DNA. Resumption of DNA synthesis can occur by replacement of the replicative DNA polymerase with specialized, error-prone translesion DNA polymerases (TLS), that have higher tolerance for damaged substrates. Several of these polymerases (Polλ, Polη and PrimPol) are stimulated in DNA synthesis through interaction with PolDIP2, however the mechanism of this PolDIP2-dependent stimulation is still unclear. Here we show that PrimPol uses a flexible loop to interact with the C-terminal ApaG-like domain of PolDIP2, and that this contact is essential for PrimPol’s enhanced processivity. PolDIP2 increases PrimPol’s primer-template and dNTP binding affinity, which concomitantly enhances PrimPol’s nucleotide incorporation efficiency. This activity is dependent on a unique arginine cluster in PolDIP2 and could be essential for PrimPol to function *in vivo*, since the polymerase activity of PrimPol alone is very limited. This mechanism, where the affinity for dNTPs gets increased by PolDIP2 binding, could be common to all other PolDIP2-interacting TLS polymerases, i.e. Polλ, Polη, Polζ and REV1, and might be critical for their *in vivo* function of tolerating DNA lesions at physiological nucleotide concentrations.

## INTRODUCTION

In all organisms, the bulk of genomic DNA replication is done by replicative DNA polymerases. These enzymes, characterized by their high processivity and fidelity, also encounter obstacles on the DNA that significantly delay progression, such as damaged DNA. If not alleviated, this can lead to fork collapse and genomic instability. In humans, genomic instability is tightly associated with disease, most notably cancer (1–3). In order to prevent genomic instability, cells have developed several DNA damage tolerance mechanisms that help the replication fork deal with various disturbances and allow completion of the replication process.

One mode of DNA damage tolerance is <u>T</u>rans<u>l</u>esion DNA <u>s</u>ynthesis (TLS), that uses specialized DNA polymerases, which at the expense of processivity and fidelity, can continue synthesis through the damaged sequence and therefore ensure the replication fork progression (4-6). Among various TLS polymerases, the primase/polymerase PrimPol has been shown to help both in nuclear and mitochondrial DNA replication fork progression (7–8). Additionally, PrimPol can also re-prime downstream of blocking lesions, thus re-reinitiating DNA replication (1, 9–15). PrimPol is a monomeric enzyme belonging to the archaeal-eukaryotic primase (AEP) superfamily (16). Its catalytic core contains three highly conserved motifs (see Figure1A) which build the dNTP binding site used for elongation (17). To start primer synthesis, PrimPol requires a unique zinc finger (ZnF)-containing C-terminal domain, which facilitates binding of the first 5’-nucleotide (1, 18). Beside its function as a primase, PrimPol behaves *in vitro* as a polymerase with very low processivity (9, 19), suggesting the need for a cofactor to function optimally in the cell. In contrast to some other TLS polymerases (20), PrimPol’s activity is not regulated by the sliding clamp Proliferating Cell Nuclear Antigen (PCNA) (21), suggesting that another factor might control its function. Indeed, polymerase δ-interacting protein 2 (PolDIP2; also known as PDIP38) and single stranded DNA binding protein Replication Protein A (RPA) (22– 24) are able to stimulate PrimPol-dependent DNA synthesis. Interestingly, PolDIP2 was previously shown to physically interact with TLS polymerases Polζ and REV1 (25), as well as to increase the processivity of Polλ and Polη (26, 27). Therefore, many TLS DNA polymerases could be regulated by a common, still to be elucidated, PolDIP2-dependent mechanism.

**Figure 1.**
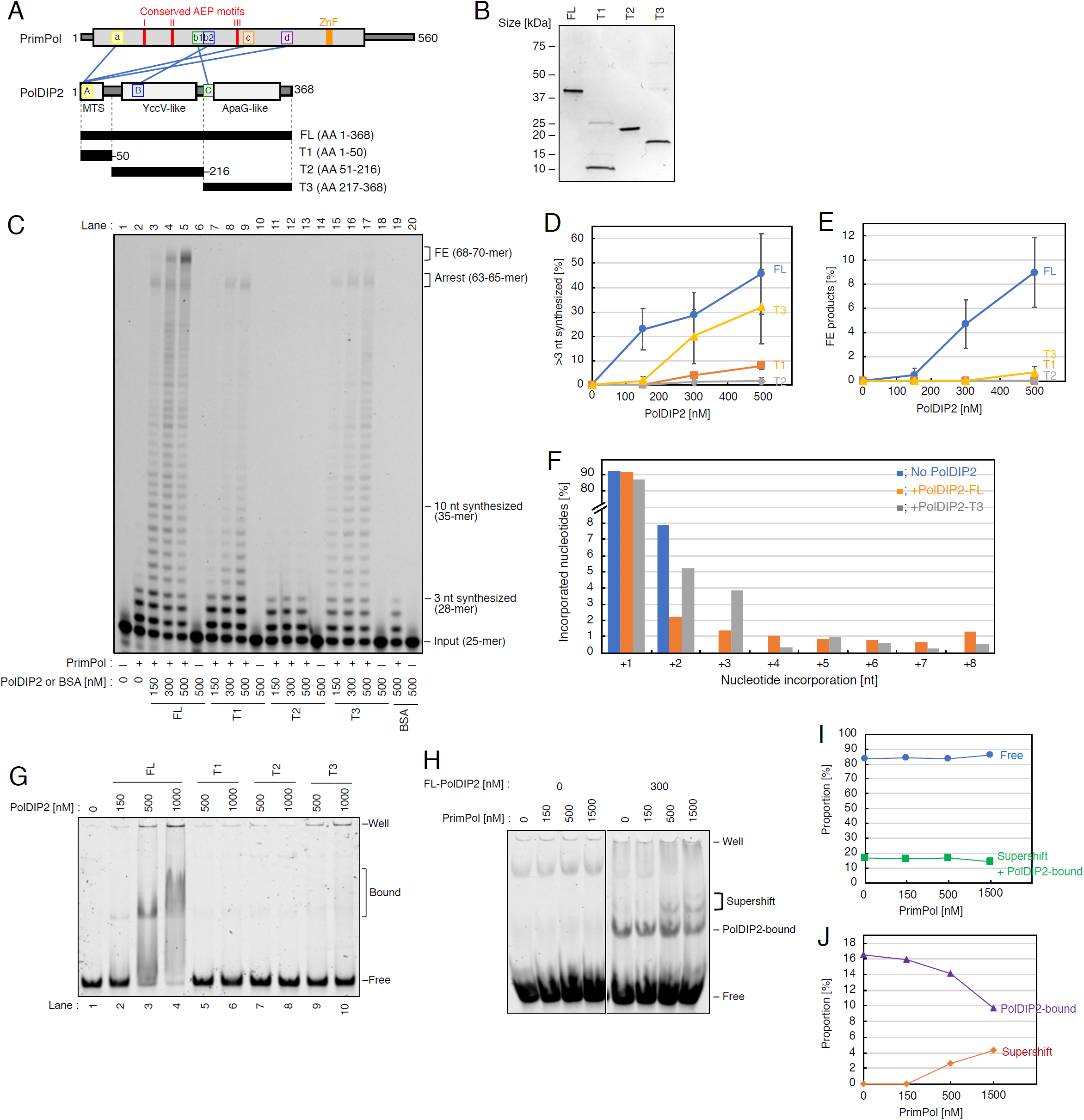
PrimPol DNA polymerase stimulation by the PolDIP2 ApaG-like domain. (A) Schematic presentation of the PrimPol-PolDIP2 interactions. The conserved AEP motifs of PrimPol (I, II and III) are indicated with red lines, and the zinc finger (ZnF) as an orange box. The PrimPol regions involved in PolDIP2 interaction are indicated as, “a”, “b1”, “b2”, “c” and “d”, and the PolDIP2 regions involved in PrimPol interaction are indicated as, “A”, “B” and “C”. Three PolDIP2 fragments, T1-, T2-, and T3-PolDIP2, harboring the mitochondrial targeting signal (MTS), YccV-like domain, and ApaG-like domain, respectively, were purified for this study. The interaction sites between PrimPol and PolDIP2, identified previously by crosslinking-mass spectrometry analysis (22), are shown with blue lines. (B) Purification of PolDIP2 fragments. 300 ng of each purified PolDIP2 fragment on SDS-PAGE gel stained with InstantBlue (Expedeon, UK). (C-E) Stimulation of PrimPol polymerase processivity by PolDIP2. (C) Reactions with 150 nM of PrimPol in the presence of 15 nM of 5’-TET-labeled primer/template DNA and the indicated amount of PolDIP2 fragments, or BSA (negative control). The positions of full extension (FE) products (68-70-mer), arrest site (63-65-mer), products extended for >10 nt (>35-mer) or for >3 nt (>28-mer), and input DNA (25-mer) are indicated. This experiment was repeated 3 times, and we quantified the percentage of (D) >3 nt synthesized and (E) full extension DNA products. The average values with standard deviations are shown with error bars. (F) Termination probabilities (y-axis) at positions between +1 and +8 under conditions where the product is derived from a single cycle DNA synthesis (single-hit conditions). The primer/template DNA (50 nM) was present at a 10-fold excess molarity over PrimPol (5 nM) to minimize the re-initiation of synthesis on the same template after an initial termination event. (G) PolDIP2-DNA binding assessed by EMSA. “Well” - gel well. “Bound” - DNA-PolDIP2 complexes. “Free” - protein-free DNA. (H-J) FL-PolDIP2 can stimulate PrimPol binding to DNA. (H) “Well” - gel well. “Supershift” - the complexes dependent on both PrimPol and PolDIP2. “PolDIP2-bound” - DNA-PolDIP2 complexes. “Free” - protein-free DNA. Quantification of Figure 1H, showing (I) the proportion of protein-free or bound DNA, and (J) DNA bound to PolDIP2 and PrimPol-dependent complexes.

Here we show that PrimPol uses a flexible loop located at its AEP catalytic core to interact with the conserved ApaG-like C-terminal region of PolDIP2 (28), which is essential for the stimulation of its primer extension activity. Using a structural model of PolDIP2 ApaG-like domain, we identify a unique arginine cluster that we show is critical for PrimPol stimulation. Moreover, nucleotide binding assays show that these arginine residues are able to assist dNTP recruitment to the PrimPol-PolDIP2 complex. Taken together, our data shows that PolDIP2 stimulates PrimPol’s processivity through stabilization of the primer-template DNA binding, but also by increasing the affinity for the incoming dNTPs. This processivity stimulation mechanism could be common to all other PolDIP2-interacting TLS polymerase such as Polλ, Polη, Polζ and REV1 and might be critical for their *in vivo* function in DNA damage tolerance, specially at physiological nucleotide concentrations.

## MATERIALS AND METHODS

### Purification of recombinant proteins

Wild type human PrimPol, as well as PrimPol Δ56 (lacking amino acids 203-258), were expressed and purified as previously described (9, 18, 19). For purification of full-length (FL) human PolDIP2, a DNA fragment encoding FL-PolDIP2 was cloned into pET21d (+) plasmid. This construct was also used for the preparation of expression vectors and purification of all other truncated or mutated PolDIP2 protein variants, except for T1-PolDIP2. Plasmids coding for FL-PolDIP2 and T2-PolDIP2 were transformed in *E. coli* ArcticExpress (DE3) cells. An overnight culture was grown at 30 °C, and used for inoculation of 1.5 L LB medium that was incubated at 30 °C until A_600_ 0.8. The temperature was then reduced to 12 °C and protein expression induced by addition of 0.4 mM IPTG. After 18 h of incubation, cells were collected by centrifugation at 4,000 × g for 15 min at 4 °C. Pellets were washed with ice-cold PBS and then resuspended in lysis buffer A (40 mM Tris-HCl pH 7.5, 5% glycerol, 500 mM NaCl, 1 mM DTT, 0.5% Tween 20, 1 μM Pepstatin, 0.1 mM AEBSF, 1 mM DTT and 1 mM E64). Cells were lysed by pulse-sonication of cell suspensions on ice, and cleared by centrifugation at 25,000 g for 30 min at 4°C. Cell lysates were supplemented with 10 mM imidazole and incubated with Ni-Sepharose excel resin (GE Healthcare) for 2 that 4 °C. The beads were washed with buffer B (40 mM Tris-HCl pH 7.5, 5% glycerol, 500 mM NaCl) with 20 mM imidazole, buffer B with 50 mM imidazole, and the proteins were eluted with buffer B containing 500 mM imidazole. Elution fractions containing PolDIP2 were concentrated using a Vivaspin 2 column, and further purified by gel filtration using Superdex 200 Increase 10/300 GL column with buffer C (40 mM Tris-HCl pH 7.5, 5% glycerol, 100 mM NaCl). Protein samples were frozen in liquid N_2_ and stored at −80 °C. For the purification of recombinant T3-PolDIP2, the expression, cell lysis and protein binding to Ni-Sepharose excel resin was performed as described for FL-PolDIP2. Before elution from the resin, bacterial chaperons were removed by incubating the protein in buffer C with 5 mM ATP and 5 mM MgCl_2_. Eluted fractions containing T3-PolDIP2 were loaded on a SP Sepharose High Performance column (GE Healthcare), using buffer D (40 mM Tris-HCl pH 7.5, 5% glycerol, 25 mM NaCl) and eluted in buffer C. To avoid aggregation, eluted fractions were diluted with the same volume of buffer C with 80% glycerol and store at −20°C. As for purification of T1-PolDIP2, a DNA fragment with GST tag and TEV protease recognition site was introduced in N-terminal region of T1-PolDIP2. Recombinant T1-PolDIP2 was first purified with Glutathione Sepharose 4 Fast Flow. The cell lysate was incubated with Glutathione Sepharose 4 Fast Flow resin for 18 h at 4 °C. The resin was washed with buffer B, and the protein was eluted with buffer B containing 40 mM Glutathione, reduced (Duchefa Biochemie). After removal of the nGST tag with TEV protease, fractions containing T1-PolDIP2 were supplemented with 10 mM imidazole and incubated with Ni-Sepharose excel resin, washed with buffer B containing 20 and 50 mM imidazole. T1-PolDIP2 was eluted with buffer B containing 500 mM imidazole. Protein samples were frozen in liquid N_2_ and stored at −80 °C.

### Oligonucleotides

For DNA replication assays and electrophoretic mobility shift assays (EMSA) in Figures 1G, 1H, and Supplementary Figure S3A, a 5’-TET fluorophore-labeled 25-mer oligonucleotide primer (5’-TET-ATAGGGGTATGCCTACTTCCAACTC-3’) was hybridized to a 70-mer oligonucleotide template (5’-GAGGGGTATGTGATGGGAGGGCTAGGATATGAGGTGAGTTGAGTGGAGTTGGAAGTAGGCATACCCCTAT-3’) (19). In filter binding assays in Figure 5, a non-labeled 25-mer primer (5’-ATAGGGGTATGCCTACTTCCAACTC-3’) was hybridized with the 70-mer oligonucleotide template. For EMSA in Supplementary Figure S3B, the 5’-end of 60-mer “GTCC” oligonucleotide (5’-T_36_CCTGT_20_-3’) was labeled with [γ-^32^P]ATP.

### Primer extension assay

The reaction mixture (10 μL) contained 10 mM Bis-Tris propane (pH 7.0), 40 mM NaCl, 10 mM MgCl_2_, 1 mM DTT, 200 *µ*M dNTPs, 15 nM of hybridized 5’-TET-primer/template DNA, indicated amounts of purified PrimPol (WT or Δ56 mutant) and PolDIP2 (WT, truncated proteins, or mutants). After incubation for 10 min at 37 °C, reactions were stopped by addition of 10 *µ*l of formamide loading buffer (0.5% SDS, 25 mM EDTA, 95% v/v formamide and xylene-cyanol). For single hit experiments shown in Figure 1F and Supplementary Figure S2, the reaction composition was modified to include 50 nM hybridized primer/template DNA, 5 nM PrimPol WT, and indicated amounts of PolDIP2 fragments. Those reaction mixtures were incubated at 37°C for 30 min. DNA products of primer extension assays were separated on 10% polyacrylamide gels containing 7 M urea, and directly imaged using a Typhoon 9400 scanner (Amersham Bioscience). The images were were quantified with ImageQuant TL 8.1 software (GE healthcare). The percentage of primer that was extended for >3nt, >10 nt or was fully extended, was calculated as the ratio between the signal of corresponding products and the signal of the 25-nt primer in the control reaction without protein. When used, the polymerisation termination probabilities at specific nucleotides were calculated as previously described (29).

### Electrophoretic mobility shift assays (EMSA)

In the experiment presented in Figure 1G, the reaction mixture (10 μL) contained 10 mM Bis-Tris propane (pH 7.0), 10 mM MgCl_2_, 1 mM DTT, 15 nM of hybridized 5’-TET-primer/template DNA and indicated amounts of purified PolDIP2 fragments (FL, T1, T2, or T3). After incubation for 5 min at 37 °C, DNA-protein complexes were separated by 10% polyacrylamide gel in Tris-borate buffer, and imaged using a Typhoon 9400 scanner. The percentage of bound DNA was calculated as the ratio between the signal of bound DNA and the signal of the DNA template in the control reaction without protein. In the experiments presented in Figure 1H and Supplementary Figure S3A, the reaction mixture (10 μL) contained 10 mM Bis-Tris propane (pH 7.0), 300 mM NaCl, 1 mM CaCl_2_, 1 mM DTT, 5 *µ*M dCTP, 15 nM of hybridized 5’-TET-primer/template DNA, and indicated amounts of PrimPol and PolDIP2. After 5 min incubation at 37 °C, DNA-protein complexes were separated by 6% polyacrylamide gel in Tris-borate buffer at 4°C, and imaged using a Typhoon 9400 scanner. In the experiment presented in Supplementary Figure S3B, the reaction mixture contained 50 mM Tris-HCl [pH 7.5], 50 mM NaCl, 1 mM MnCl_2_, 1 mM DTT, 2.5% glycerol, 0.1 mg/ml BSA and 2.5 nM polyethylenglycol 4000, 2.5 nM 5’-[γ-^32^P]-labeled 60-mer “GTCC” oligonucleotide, and indicated concentrations of PrimPol or FL-PolDIP2. After 20 min incubation at 25°C, DNA-protein complexes were separated by 6% polyacrylamide gel in 1x Tris-glycine buffer at 4°C. The gels were then dried and imaged by autoradiography using a Typhoon 9400 scanner.

### Filter-based PrimPol-dNTP binding assay

The reaction mixture (10 μL) contained 10 mM Bis-Tris propane (pH 7.0), 10 mM CaCl_2_, 1 mM DTT, 1 *µ*M dCTP including [α-^32^P]dCTP, 300 nM of hybridized primer/template DNA and 600 nM of purified human PrimPol WT or PolDIP2-FL (WT or mutants). After incubation for 5 min at 37 °C, the reaction mixture was loaded on a nitrocellulose membrane (0.45 *µ*m; Amersham), and washed with 1 ml of washing buffer (10 mM Bis-Tris propane (pH 7.0), 10 mM CaCl_2_, and 1 mM DTT). The membrane was transferred to a tube and the protein-bound [α-^32^P]dCTP was quantified in a Beckman Coulter LS 6500 Liquid Scintillation Counter using OptiPhase HiSafe 3 (PerkinElmer) as solvent.

### Structural comparisons and computational simulation of PolDIP2

Multiple sequence alignments of different PrimPol species, or human PolDIP2 versus human Fbxo3 and bacterial ApaGs, were performed using COBALT (constraint-based multiple alignment tool (30, 31) from the National Center for Biotechnology Information (NCBI). The homology models of T3-PolDIP2 (amino acids (AA) 217-368) WT, 3R>A, and R282 were constructed based on the crystal structure of *Shewanella oneidensis* ApaG protein (PDB-ID 1TZA), using I-TASSER web service. Three-dimensional images and surface electrostatic potential calculation were carried out using the Swiss-PdbViewer online server application (http://www.expasy.org/spdbv/) (32).

## RESULTS

### The C-terminal region of PolDIP2 increases the processivity of PrimPol

An earlier study showed that human PolDIP2 increases the DNA polymerase activity of PrimPol (22). The crosslinking experiments in that study identified several regions as potential PrimPol-PolDIP2 interaction sites: however, the mechanism behind the PolDIP2-dependent stimulation of PrimPol DNA synthesis remained unclear. To understand how this PrimPol stimulation is mechanistically achieved, we purified four cHis6-tagged PolDIP2 fragments (Figure 1A and 1B). Apart from the FL-PolDIP2 protein (AA 1-368), we purified an N-terminal T1 fragment (AA 1-50) which covers the mitochondrial targeting sequence (MTS), a middle T2 fragment (AA 51-216) which contains the YccV-like domain, and a C-terminal T3 fragment (AA 217-368) which contains an ApaG-like domain (Figure 1A and 1B) (28). In addition to representing distinct functional parts of the protein, the fragments also include the three different PrimPol-interaction regions within PolDIP2 (Figure 1A) (22).

To determine if any of these isolated domains of PolDIP2 could sustain the stimulation of PrimPol DNA polymerization, we performed primer-extension assays on a 25/70-mer DNA primer/template substrate in the presence of increasing concentration of each PolDIP2 variant (Figure 1C). PrimPol alone showed limited processivity under the set reaction conditions, with the 25 nt 5’-labelled primer extended by 1 to 3 nt (Figure 1C, lane 2). Addition of FL-PolDIP2 stimulated DNA synthesis by PrimPol, resulting in long DNA products (46 % of the DNA substrate was extended further than 3 nt in the presence of 500 nM FL-PolDIP2; Figure 1D), which included run-off products that are 70 nt in length (Figure 1C lanes 3-5; Figure 1E). This result is consistent with previous work and confirms that FL-PolDIP2 is able to stimulate PrimPol during DNA synthesis (22). Primer-extension reactions in the presence of the PolDIP2 fragments showed the PrimPol stimulation is mainly dependent on fragment T3 (Figure 1C, lanes 15-17) and to a lesser extent on T1 (Figure 1C, lanes 7-9), but not on T2 (Figure 1C, lanes 11-13). Even the lowest concentration of T3-PolDIP2 fragment (150 nM; Figure 1C, lane 15) showed a robust stimulation of DNA synthesis. Nonetheless, a small reduction of the >28 nt DNA products were detected when compared to the reaction with FL-PolDIP2 (Figure 1C, compare lanes 5 and 17; 32% T3 *vs* 46% FL in Figure 1D), and fully extended DNA products (68-70 nt) were not accumulated in the presence of T3 at any concentration (Figure 1C and 1E). Interestingly, addition of 300-500 nM T1-PolDIP2 also showed a modest increase in PrimPol DNA synthesis (Figure 1C, compare lane 2 with lanes 8 and 9; Figure 1D), but this T1 stimulation was substantially weaker compared to the effect of T3- or FL-PolDIP2 (Figure 1C, compare lanes 7-9 with lane 15-17; Figure 1D).

We also purified the putative mitochondrial variant of the protein (AA 51-368), comprising both domains T2 and T3 (Supplementary Figure S1A and S1B). Consistent with the previous study (22), this mitochondrial variant of PolDIP2 (Fragments T2+T3) was unable to stimulate PrimPol DNA synthesis (Supplementary Figure S1C).

To exclude that PolDIP2 affects DNA product length by increasing PrimPol re-loading efficiency, we performed primer-extension assays with a large excess of template-primer over DNA polymerase (Figure 1F and Supplementary Figure S2). In these conditions, once a primer is extended, the probability that it will be used a second time after dissociation is negligible, and the elongated products therefore derive from a single cycle of synthesis. As observed in standard reaction conditions, both FL- and T3-PolDIP2 considerably increased the length of products, confirming the observed stimulation comes from enhanced processivity and not PrimPol re-loading (Supplementary Figure S2). Additionally, band intensities were measured to calculate termination probability at specific nucleotide positions, as previously described (29). The presence of both FL- and T3-PolDIP2 decreased the termination probability of PrimPol synthesis at position +2; instead, longer products were also observed (Figure 1F (positions +3 to +8) and Supplementary Figure S2), indicating a gain in PrimPol’s processivity during DNA polymerization.

EMSA experiments showed that FL-PolDIP2 is able to bind to the primer-template substrate used in the primer extension assays (Figure 1G, lanes 3 and 4), but that separate fragments (T1, T2 and T3) show little or no DNA binding (lanes 5-10). PrimPol has no detectable stable binding to the primer-template substrate by itself (Figure 1H); however, FL-PolDIP2 is able to enhance the primer-template DNA binding affinity of PrimPol, as demonstrated by the appearance of a supershifted band in the presence of the two proteins (Figure 1H, 1I and 1J). On the contrary, T3-PolDIP2 was unable to alter PrimPol’s template-primer binding (Supplementary Figure S3A). Thus, the stimulation of the polymerase activity of PrimPol by the T3 fragment seems to be independent on the intrinsic DNA binding capacity. PrimPol and FL-PolDIP2 are both able to bind to PrimPol’s preferred 60-mer GTCC ssDNA substrate, but we were unable to demonstrate a collaboration of the two proteins in ssDNA binding (Supplementary Figure S3B). This might indicate that FL-PolDIP2 stabilizes PrimPol’s binding to the 3’ end of the primer, but does not alter its general affinity for the DNA template.

In conclusion, whereas the interaction of PrimPol and FL-PolDIP2 stabilizes PrimPol’s binding to a primer-template substrate, thus explaining an increase in processive polymerization, the stimulation in processivity observed with the T3 fragment of PolDIP2 does not correlate with an improvement in DNA binding.

### A flexible loop in the PrimPol DNA polymerase core is essential for PolDIP2-dependent stimulation

As shown here, the most important PolDIP2 fragment for enhancing PrimPol processivity is T3. Within this fragment, a region here termed “C” (AA 215-223), has been previously shown to directly interact with region “b1” (AA 226-232) in PrimPol (Figure 1A, 2A and 2B) (22). These PrimPol residues are located within a non-structured, perhaps flexible loop (17), spanning AA 201 to 260 of human PrimPol (Figure 2A and 2B). This disordered region is relatively conserved in primates, but largely diverged in PrimPols from other vertebrates and plants; however, the signature of the PolDIP2 interacting region “b1” is still present in PrimPols from rodents and amphibians (Figure 2A).

**Figure 2.**
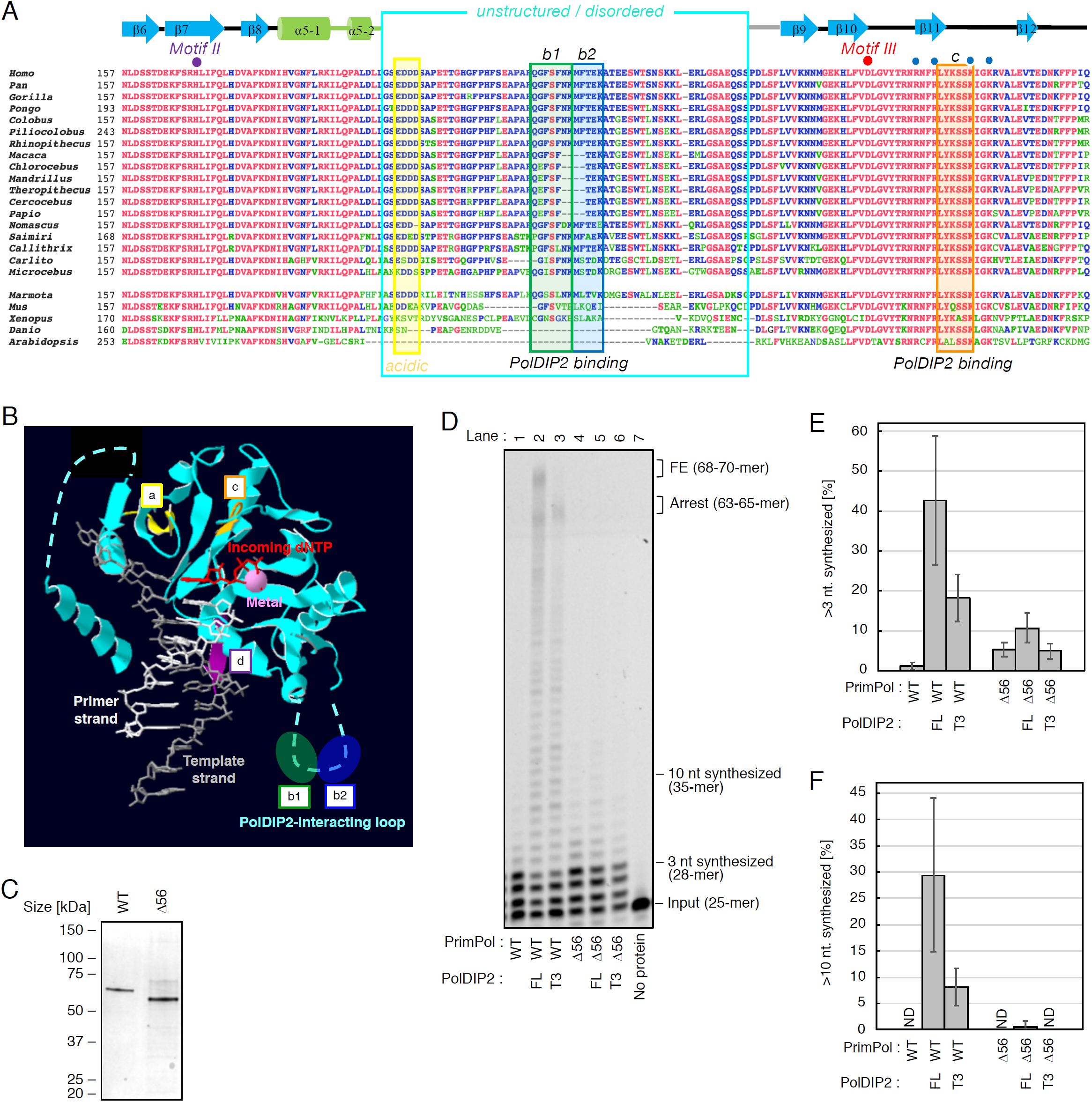
PrimPol AA 203-258 are required for PolDIP2-depended stimulation of DNA synthesis. (A) Multiple amino acid sequence alignment of different PrimPol species, in the region spanning AEP motifs II and III. PolDIP2 interacting regions “b1”, “b2” and “c” are framed in green, blue, and orange, respectively. The unstructured/disordered loop region, deleted in the Δ56 (AA 203-258) PrimPol mutant, is indicated in cyan. The secondary structure elements of human PrimPol (beta-strands and alpha-helices) are indicated above the human sequence. Individual residues involved in dNTP binding (purple dot), catalysis (red dot) or phosphate binding (blue dots) are indicated. The references of the aligned sequences are: *Homo sapiens* (NP_689896); *Pan troglodytes* (XP_001162592); *Gorilla gorilla* (XP_004040738); *Pongo abelii* (XP_024101228); *Colobus angolensis* (XP_011814937); *Piliocolobus tephrosceles* (XP_023076725); *Rhinopithecus roxellana* (XP_010376227); *Macaca fascicularis* (XP_005556457); *Chlorocebus sabaeus* (XP_007998585); *Mandrillus leucophaeus* (XP_011855552); *Theropithecus gelada* (XP_025240422); *Cercocebus atys* (XP_011941669); *Papio anubis* (XP_003899457); *Nomascus leucogenys* (XP_003271526); *Saimiri boliviensis* (XP_010342695); *Callithrix jaccus* (XP_008990643); *Carlito syrichta* (XP_008055415); *Microcebus murinus* (XP_012640401); *Marmota marmota* (XP_015341649); *Mus musculus* (XP_017168376); *Xenopus laevis* (XP_018086024); *Danio rerio* (NP_001032455); *Arabidopsis thaliana* (NP_001154775). (B) The reported human PrimPol structure, with a primer/template DNA, metal and incoming dNTP (17), has a non-structured flexible loop (containing sites “b1” and “b2”), in addition to sites “a”, “c” and “d”, that interact with PolDIP2 (22). (C) 300 ng of purified WT PrimPol or Δ56 PrimPol mutant on SDS-PAGE gel stained with InstantBlue. (D) Primer extension reactions using 150 nM of PrimPol WT or Δ56 incubated with equimolar amounts (150 nM) of FL-PolDIP2 or T3-PolDIP2, as indicated. These experiments were performed twice, and we quantified the percentage of (E) >3 nt and (F) >10 nt synthesized DNA products. The average values with standard deviations are shown.

To investigate if this PrimPol-PolDIP2 interaction is required for PrimPol stimulation, we purified a PrimPol variant (Δ56) with deleted flexible loop (lacking AA 203-258; Figure 2C). Whereas the WT and Δ56 PrimPol variants have comparable primer extension activities, showing that the flexible loop is dispensable for PrimPol’s DNA polymerization (Figure 2D, compare lanes 1 and 4), only the WT PrimPol was stimulated by FL- and T3-PolDIP2 (Figure 2D, lanes 2-3; quantitative analysis in Figure 2E). On the contrary, both were unable to efficiently stimulate the Δ56 PrimPol variant (Figure 2D, lanes 5-6; Figure 2E). Addition of FL-PolDIP2 still shows a minimal stimulation of Δ56 PrimPol (Figure 2D, compare lanes 4 and 5; Supplementary Figure S4A-C), as judged by quantitation of both short (Figure 2E) and long (Figure 2F) extension products, whereas the T3-PolDIP2 fragment was completely unable to stimulate the Δ56 PrimPol variant (Figure 2D, lane 6; Figure 2E and 2F), even when higher concentrations of the T3 variant were used (Supplementary Figure S4A-C). In conclusion, the flexible loop in PrimPol (AA 201-260), which contains a binding site for the interaction with the T3 region, is essential for PolDIP2-dependent stimulation of PrimPol DNA polymerization.

### PolDIP2 contains a putative pyrophosphate binding motif and an Arginine cluster

Sequence comparison shows that the T3-PolDIP2 fragment contains a bacterial protein ApaG-like domain that also shows sequence similarity with the human F-Box only 3 (FBxo3) protein. A multiple amino acid sequence alignment of the T3 domain of human PolDIP2 with human FBxo3 and the closest ApaGs from various species confirmed the structural similarity (Figure 3A). Strikingly, some of these aligned sequences also contain the region “C” of PolDIP2, involved in interaction with PrimPol (16; cyan box in Figure 3A). More importantly, one of the most conserved portions of this alignment, present in all three types of proteins, shares a loop with a highly conserved motif (GxGxxG), that has been proposed to be involved in binding to the pyrophosphate moiety of a nucleotide derivative (33, 34). The crystal structure of *S. oneidensis* ApaG (PDB-ID 1TZA) (35) shows that the putative pyrophosphate binding motif (G^65^xG^67^xxG^70^) forms a double turn between two beta strands (β4 and β5; see Figure 3A) that builds a wall at the end of a channel that serves a cavity to bind pyrophosphate (Figure 3B, left). At the bottom of this channel, a specific arginine (Arg^49^) appears to be the likely candidate to interact with the phosphate moiety of nucleotide derivatives. As the amino acid sequence identity between T3-PolDIP2 and *S. oneidensis* ApaG was about 30%, high enough to generate a homology model (36), we built a structural model of the T3-PolDIP2 fragment (Figure 3B, right; Supplementary Figure S5A and S5B). Interestingly, the T3 domain of PolDIP2 also contains an apparent binding channel, but with a unique positively-charged Arginine cluster (Arg^282^, Arg^297^, Arg^299^) at the bottom, that partly overlaps with its putative pyrophosphate binding motif (298-<u>G</u>R<u>G</u>VV<u>G</u>) (Figure 3A and 3B). The first of these Arginine (Arg^282^) residues aligns with Arg^49^ of *S. oneidensis* ApaG, whereas the other two residues (Arg^297^, Arg^299^) are highly conserved in PolDIP2 of higher eukaryotes, and occasionally in some ApaG sequences (Figure 3A).

**Figure 3.**
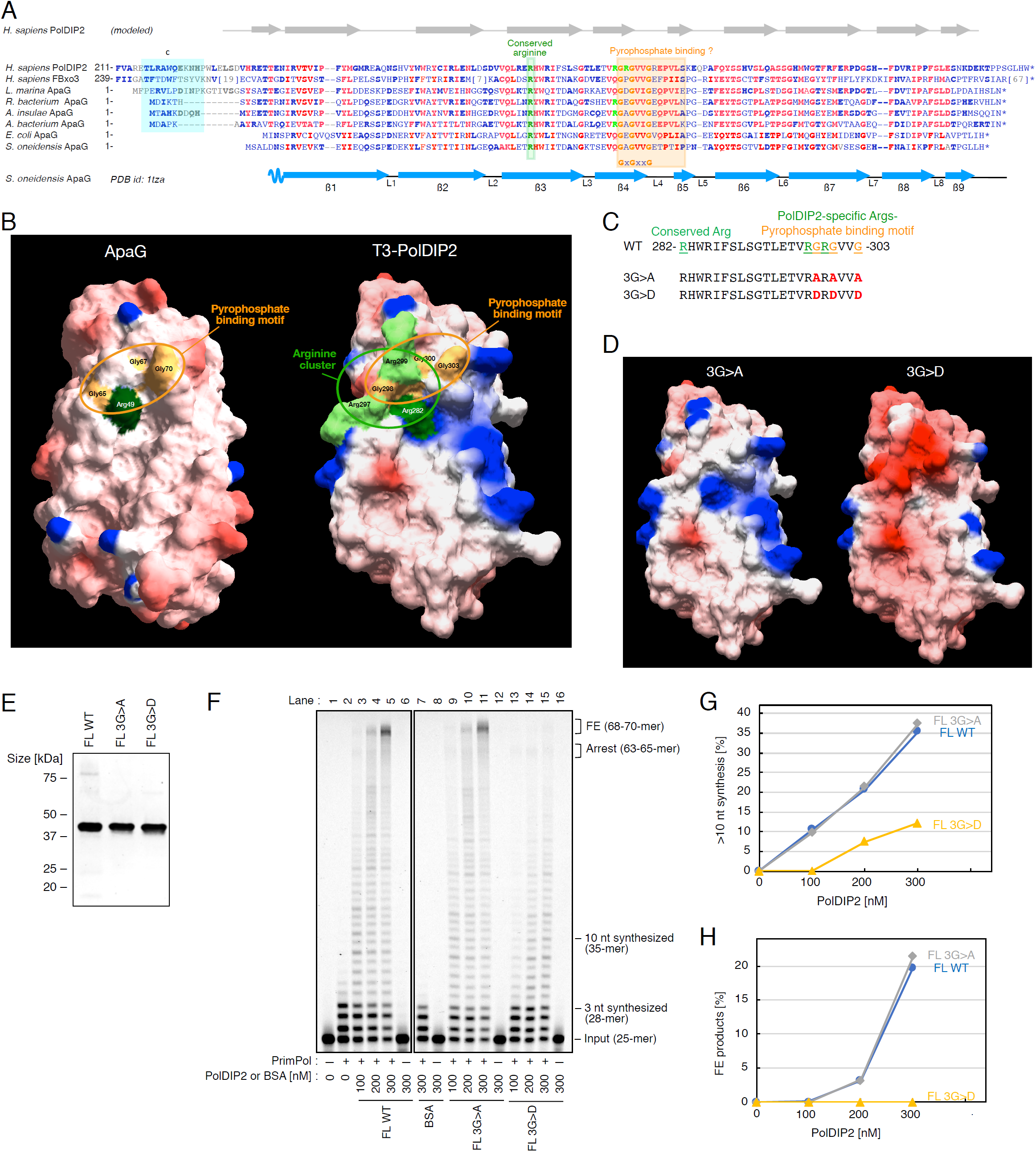
The conserved PolDIP2 pyrophosphate binding motif is dispensable for PrimPol stimulation. (A) The T3 domain of human PolDIP2 (AA 217-368) has homology to FBxo3 and ApaG proteins, and contains a pyrophosphate binding motif and a proximal Arg-cluster. The amino acid sequences aligned were: *Homo sapiens* PolDIP2 (NP_056399); *Homo sapiens* FBxo3 (XP_011518282.1); *Labrenzia marina* ApaG (POF34899.1); *Rhodospirillaceae bacterium* ApaG (HBC05999.1); *Aestuariispira insulae* ApaG (RED46168.1); *Alphaproteobacteria bacterium* ApaG (OLB70390.1); *Escherichia coli* ApaG (ART45711.1); *Shewanella oneidensis MR-1* ApaG (AAN56626). Region “C” in PolDIP2, involved in interaction with the “b1” region of PrimPol is indicated with a cyan box. The positions of the pyrophosphate binding motif (boxed in orange), G^298^xG^300^xxG^303^ in human PolDIP2, and the Arg-cluster residues Arg^282^, Arg^297^, and Arg^299^ (green letters) are indicated. Residues with the highest conservation are indicated in red, and other conserved residues are indicated in bold blue letters. The secondary structure elements (beta-strands β1 to β9 and loops L1 to L8) corresponding to *S. oneidensis* MR-1 ApaG (PDB id: 1TZA) is indicated below the multiple alignment. The secondary structure elements of the predicted model for the T3 domain of *H. sapiens* PolDIP2 are indicated above the multiple alignment. (B) Left: surface electrostatic potential of the *S. oneidensis* MR-1 ApaG (PDB id 1TZA), Right: surface electrostatic potential of the 3D-structure model for T3-PolDIP2, obtained using the I-TASSER web tool with PDB-ID 1TZA as a template. The residues of the Arg-cluster and pyrophosphate binding motif are shown with green and orange, respectively. The amino acid sequence (C) and the structural models (D) of PolDIP2 pyrophosphate binding motif mutants used in this study. Two triple mutations 3G>A (G298A/G300A/G303A) and 3G>D (G298D/G300D/G303D) were introduced into FL-PolDIP2. (E) 300 ng of purified FL-PolDIP2 WT or mutants (3G>A or 3G>D) on SDS-PAGE gel stained with InstantBlue. (F) Primer extension experiments using 150 nM of WT PrimPol in the presence of the indicated amounts of FL-PolDIP2 WT, 3G>A, or 3G>D. The ratio of DNA products extended for >10 nt (G), or fully extended (H).

### The Arginine cluster in PolDIP2 is essential for stimulation of PrimPol DNA synthesis

To investigate if the pyrophosphate binding motif in PolDIP2 has an essential role in PrimPol stimulation, we purified two mutant variants of FL-PolDIP2 with alterations in the G^298^xG^300^xxG^303^ motif. A triple mutant conservative replacement 3G>A (G298A/G300A/G303A) which was predicted to moderately affect the functionality of the pyrophosphate binding motif, and a more radical triple substitution 3G>D (G298D/G300D/G303D) that was expected to dismantle the double turn connecting beta strands, and perhaps also affect the surrounding electrostatic potential (Figure 3C-E). Our results show that the conservative alteration of the pyrophosphate binding motif did not affect PrimPol stimulation, since the FL-3G>A mutant had a wild-type-like activity, stimulating PrimPol in primer extension assays (Figure 3F, compare lanes 3-5 with lanes 9-11; see quantitation at Figure 3G and 3H). On the other hand, the Gly-to-Asp triple mutant (FL-3G>D) shows a strong reduction in the level of PrimPol stimulation (Figure 3F, compare lanes 3-5 to lane 13-15), especially evident for primers that are extended by >10 nt (Figure 3G), or as measured by quantification of the 68-70 nts long full extention products (Figure 3H). These results suggest that the pyrophosphate binding motif is not vital for PrimPol stimulation, but on the other hand, the region surrounding the GxGxxG motif is essential, leading us to speculate that the positively charged Arg-cluster might have an important function.

To address the function of the PolDIP2 Arg-cluster in PrimPol stimulation, we purified FL-PolDIP2 variants with modifications in the Arg-cluster: a triple mutant R282A/R297A/R299A (FL-3R>A), and the single amino acid residue alterations FL-R282A, FL-R297A, and FL-R299A (Figure 4A and 4B). When tested for their ability to stimulate PrimPol in primer extension assays (Figure 4C), the FL-R299A PolDIP2 mutant displayed a FL-WT PolDIP2-like ability to stimulate PrimPol, showing that the Arg^299^ residue is not critical in this respect (Figure 4C, compare lanes 3 and 7; Figure 4D and 4E). The other single mutant variants of PolDIP2, FL-R282A and FL-R297A, showed however only a moderate capacity to stimulate PrimPol (Figure 4C, lanes 5 and 6), exposing the importance of these two residues. In agreement with these results, the triple mutant 3R>A also showed a similar reduction in the capacity to stimulate PrimPol (Figure 4C, lane 4).

**Figure 4.**
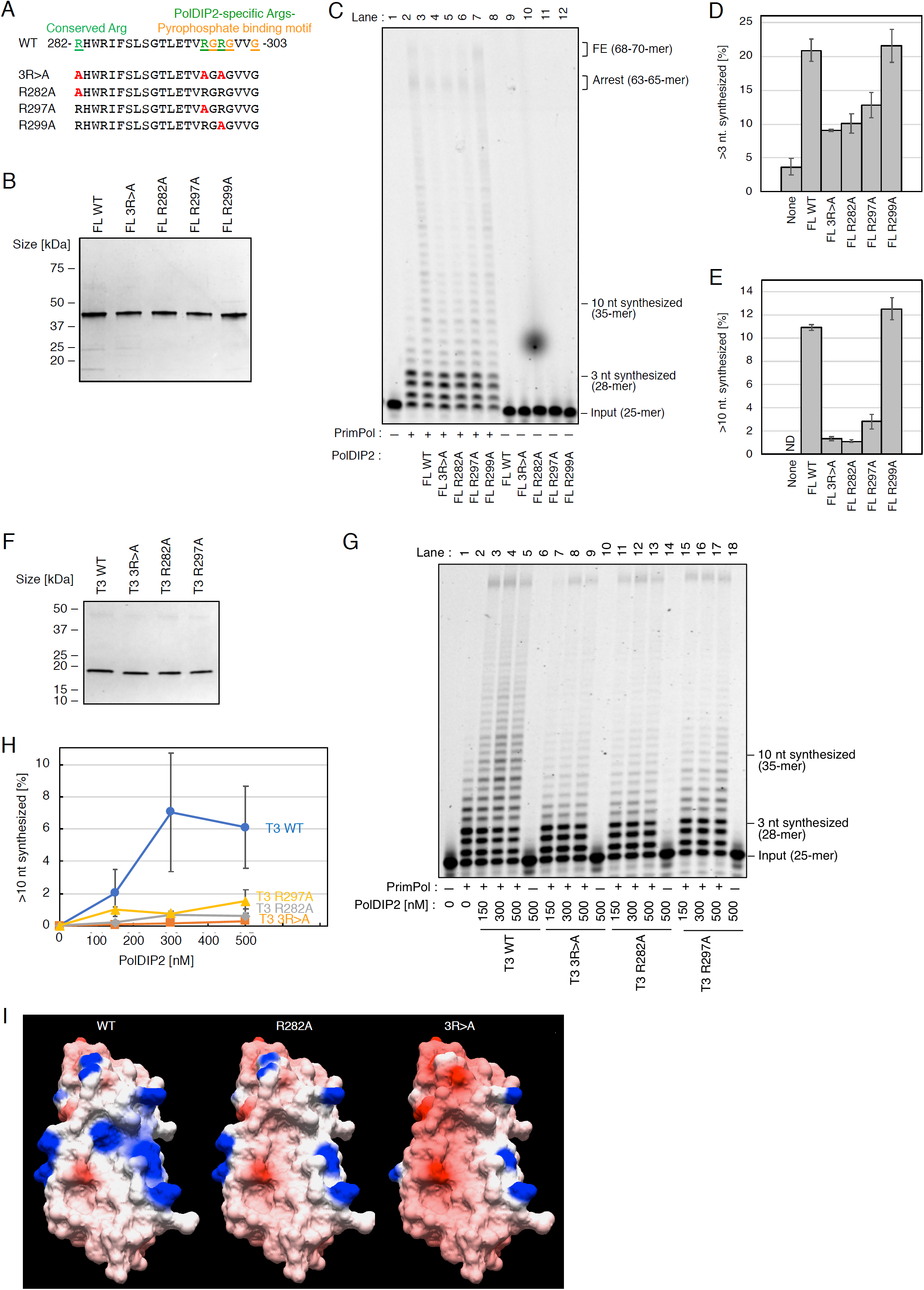
An Arg-cluster in PolDIP2 is required for PrimPol stimulation. (A) Amino acid sequences (AA 282-303) of the PolDIP2 Arg-cluster mutants used in this study. A triple mutation 3R>A (R282A/R297A/R299A), and three single Arg alterations (R282A, R297A, or R299A) were introduced into FL- or T3-PolDIP2. (B) 300 ng of purified FL-PolDIP2 WT and mutants were loaded on SDS-PAGE gel and stained with InstantBlue (C-E) PrimPol stimulation of DNA synthesis by the Arg-cluster FL-PolDIP2 mutants. (C) Primer extension reaction using 150 nM of WT PrimPol protein in the presence of equimolar amounts (150 nM) of the indicated FL-PolDIP2 variants. This experiment was repeated twice, and the percentage of (D) >3 nt and (E) >10 nt synthesized DNA products was quantified. The average values and absolute errors are shown with error bars. (F) 300 ng of purified T3-PolDIP2 WT and mutants (3R>A, R282A, or R297A) on SDS-PAGE gel stained with InstantBlue. (G-H) Stimulation of PrimPol DNA synthesis by the T3-PolDIP2 mutants. (G) Primer extension reactions with 150 nM of WT PrimPol, in the presence of the indicated amounts of T3-PolDIP2 variants. This experiment was repeated twice, and the percentage of >10 nt synthesized DNA products was quantified (H). The average values with absolute errors are shown. (I) The surface electrostatic potential of the T3-PolDIP2 variants (WT-T3, T3-R282A, and T3-R282A/R297A/R299A).

We speculated that the remaining PrimPol stimulation activity observed with FL-R282A and FL-R297A PolDIP2 mutants, clearly observed at high PolDIP2 concentrations (Supplementary Figure S6A-C) was a consequence of the moderate ability of the PolDIP2 fragment T1 to stimulate PrimPol (Figure 1C). To test this, we made the same mutant variants in the T3-PolDIP2 fragment (T3-R282A, T3-R297A and the triple mutant T3-3R>A) (Figure 4F) and found that these mutants do not stimulate PrimPol, whereas the T3-WT PolDIP2 fragment was able to increase PrimPol DNA synthesis (Figure 4G and 4H). The structural model of the T3 fragment arginine mutants (R282A and R297A) predicts a strong decrease in the positive surface charge observed in the WT PrimPol T3 fragment structural model (Figure 4I), in the vicinity of the pyrophosphate binding motif. Together these data shows that the PolDIP2 residues Arg^282^ and Arg^297^ are critical for PrimPol stimulation of DNA synthesis.

### The Arginine cluster in PolDIP2 favors dNTP binding by PrimPol

Despite the identification of critical residues in PolDIP2 and interacting regions in PrimPol, it remains unclear what is the mechanism by which T3-PolDIP2 stimulates DNA synthesis by PrimPol. One strategy to enhance the processivity of a DNA polymerase could involve increasing its binding affinity for dNTPs (37). This is particularly relevant for PrimPol, whose low affinity for dNTPs might hinder its ability to synthesize DNA *in vivo* (38, 39). To investigate if the presence of PolDIP2 influences dNTP binding by PrimPol, we performed filter binding assays in the presence of a DNA primer/template and the complementary nucleotide for the first position after the primer, [α-^32^P]dCTP. Ca^2+^ was used as the metal cofactor, since it supports PrimPol binding to the incoming dNTP, but not primer extension (Figure 5A) (17, 40). As expected, the dNTP binding capacity of PrimPol alone was much lower when compared with Pol γ, the replicative mitochondrial DNA polymerase (compare the y-axis scale of Figure 5A and 5B), and PolDIP2 alone was very inefficient in binding dCTP (Figure 5A). However, when both proteins PrimPol and PolDIP2 were added simultaneously, their likely interaction promotes a 3-fold increase in the amount of bound dCTP, compared to PrimPol alone, whereas Polγ was unaffected in dCTP binding by PolDIP2 addition (Figure 5A and 5B). Interestingly, PolDIP2 was unable to stimulate dCTP binding efficiency on ice (Figure 5A, blue bar), suggesting that a conformational change in PrimPol or/and PolDIP2 might be behind the enhanced nucleotide binding. Next, we investigated if the Arg-cluster in PolDIP2 that is critical for stimulation of PrimPol DNA polymerization, is also essential to increase dNTP binding efficiency. The PolDIP2 arginine mutants (FL-R282A, FL-R297A and FL-3R>A) and pyrophosphate binding alteration (FL-3G>D) that were unable to stimulate PrimPol DNA synthesis (Figure 3F and 4G), were also unable to increase PrimPol’s dNTP binding efficiency (Figure 5C). Conversely, PolDIP2 alterations FL-R299A and FL-3G>D that are proficient in stimulation of both PrimPol DNA synthesis (Figure 3F and 4C) are also able to stimulate the PrimPol dNTP binding efficiency (Figure 5C).

**Figure 5.**
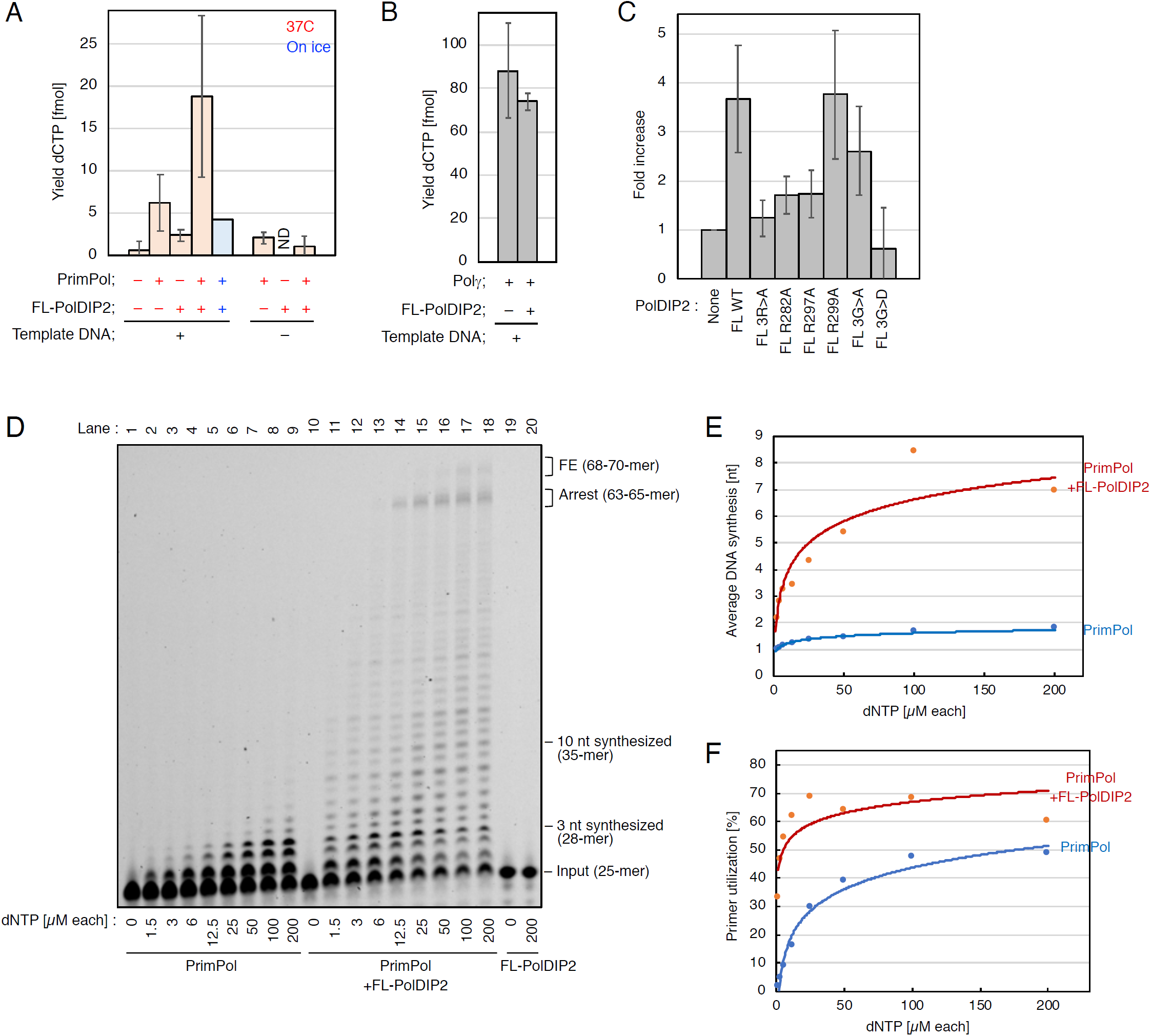
The PolDIP2 Arg-cluster increases PrimPol-dependent dNTP binding, facilitating PrimPol DNA synthesis stimulation at *in vivo* relevant dNTP levels. (A-C) Influence of PolDIP2 on the binding of dCTP to either PrimPol or Polγ. (A) dNTP binding assay was performed in the presence of 1 *µ*M dCTP including [α-^32^P]dCTP, 300 nM of primer/template DNA, 600 nM of WT PrimPol, and/or equimolar (600 nM) of WT FL-PolDIP2 incubated at 37 °C (red bars) or on ice (blue bar) for 5 min in the presence of Ca^2+^. (B) dNTP binding as described for Figure 5A, but in the presence of 300 nM PolγA, 450 nM PolγB, and 600 nM WT FL-PolDIP2. (C) The influence of the PolDIP2 Arg-cluster on dNTP binding efficiency. Conditions were as described for Figure 5A in the presence of 600 nM of the indicated FL-PolDIP2 variants. (D-F) PolDIP2-dependent PrimPol stimulation at a variety of dNTP concentrations. (D) Primer extension reactions with 150 nM of PrimPol, and 150 nM of FL-PolDIP2 in the presence of indicated dNTP concentrations. The average length of the DNA products (E) and the percentage of DNA substrates extended with 1 nt or more (F) were quantified.

A plausible hypothesis is that PolDIP2 interaction could lead to a conformational change in PrimPol that increases dNTP affinity, and that the consequent catalytic improvement would be of particular benefit at *in vivo* nucleotide concentrations (41), which are lower than those used in the experiments above. To investigate this option, we performed primer extension assays in the presence of a range of dNTP concentrations (Figure 5D). As expected, PrimPol alone poorly elongates the primer beyond nucleotide +1 at low dNTP concentrations that are thought to be physiologically relevant (up to 12,5 μM, Figure 5D, lanes 2-5). However, addition of PolDIP2 results in a substantial amount of DNA fragments extended for more than 3 nucleotides (Figure 5D, lanes 11-14). In fact, The PolDIP2-dependent stimulation of PrimPol synthesis was already evident using the lowest concentration of dNTPs (1,5 μM; Figure 5D, compare lanes 2 and 11; Figure 5E). Interestingly, PrimPol’s improved affinity for dNTPs also leads to a strong increase in primer utilization at low dNTP concentrations; the latter was only slightly increased at higher dNTP concentrations (100-200 μM; Figure 5F). Together, these data suggest that PolDIP2 stimulation of PrimPol DNA polymerase activity would be most beneficial at physiologically-relevant dNTP concentrations.

## DISCUSSION

PolDIP2 has the ability to increase the DNA synthesis efficiency of TLS DNA polymerases Polη, Polλ (19, 20), and PrimPol (22). However, previous studies have not addressed the mechanism behind the PolDIP2-dependent stimulation of DNA polymerase activity. Based on the analysis of protein-protein interaction sites between PrimPol and PolDIP2 (22), we designed several constructs and mutants of these two proteins, which allowed us to identify the critical regions that are essential for polymerization stimulation (Figure 1).

As previously reported, only the FL-PolDIP2, but not the mitochondrial variant (Mito-PolDIP2), is able to stimulate the DNA polymerase activity of PrimPol (Fig 1C, Supplementary Figure S1) (22). Because Mito-PolDIP2 lacks fragment T1, we initially suspected that the T1 segment of PolDIP2 (which interacts with three separate regions “a”, “c”, and “d” in PrimPol) might be essential for PrimPol stimulation. However, as shown in Figure 1C, the T1-PolDIP2 fragment by itself had only a minor ability to stimulate PrimPol; instead, the C-terminal region (T3) of PolDIP2 showed intrinsic and robust PrimPol stimulation. Same was observed in single turnover experiments where both FL-PolDIP2 and T3-PolDIP2 were able to increase PrimPol processivity, generating DNA products of longer length (Figure 1F). This suggests that the major contribution of PrimPol stimulation lies within the T3 domain of PolDIP2, whereas the T1 region only has a minor impact. Our results are therefore consistent with the conclusion that in Mito-PolDIP2 (i.e. fragments T2 + T3), the fragment T2 prevents fragment T3 to bind PrimPol in a productive way, thus inhibiting the stimulation observed with either full-length protein or only the T3 domain. It is at this point exciting to speculate that in mitochondria, the Mito-PolDIP2 is inactive for PrimPol stimulation. Unless other regulatory factors are able to deter the T2 fragment-dependent inhibition of T3 activity, this could indicate that PrimPol is mainly active as a primase in mitochondria (11, 15). However, this hypothesis will need to be tested in future experiments.

PolDIP2 has been proposed to increase the performance of TLS DNA polymerases in DNA damage tolerance (42). Earlier studies suggest that PolDIP2 achieves this by increasing the DNA binding affinity of TLS DNA polymerases (22, 26). Our EMSA experiments show that PolDIP2 is unable to stimulate PrimPol ssDNA binding, even at high PolDIP2 concentrations (Supplementary Figure S3B). FL-PolDIP2 is however able to increase PrimPol binding to a primer-template DNA substrate (Figure 1H), conceivably by specific stabilization of the PrimPol interaction with the 3’ terminus of the primer. The inability of the T3-PolDIP2 fragment to assist PrimPol binding to a primer-template DNA substrate (Supplementary Figure S3A) argues against T3 PrimPol stimulation through interaction of PolDIP2 with the template DNA. In agreement with this, the T3-PolDIP2 fragment shows robust PrimPol stimulation (Figure 1C) and is unable to bind primer-template DNA (Figure 1G). Instead, the stimulation requires a direct and very specific T3-PolDIP2:PrimPol interaction (Fig 2), involving PrimPol AA 201-260. This region forms a disordered loop dispensable for intrinsic DNA polymerization, but essential for PolDIP2-dependent stimulation of PrimPol, suggesting that this flexible loop could serve just as a scaffold to recruit PolDIP2. It could also be that the interaction of the T3 domain of PolDIP2 with the “b1” region embedded in the flexible loop of PrimPol promotes a much tighter interaction with the primer (Figure 6). No direct contacts with the primer strand were seen in the crystal structure of human PrimPol in ternary complex with DNA and dNTP (17). The proximity of the disordered region to the primer strand supports the possibility of a T3-mediated conformational alteration of the flexible loop resulting in an improvement of PrimPol’s primer grip. Such a reinforced binding of the primer could enhance processivity, an effect observed in PrimPol stimulation by PolDIP2.

**Figure 6.**
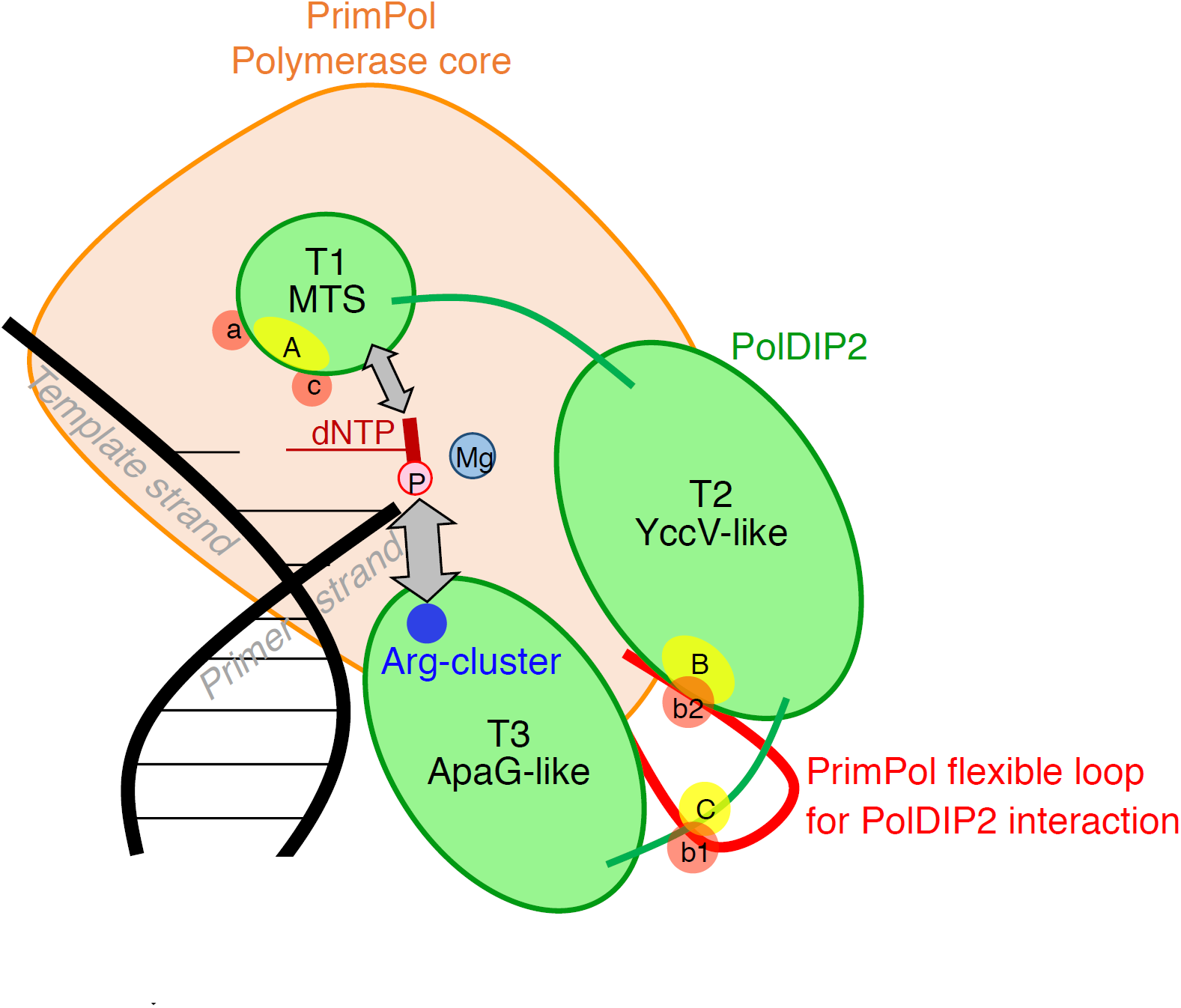
Mechanistic model for PrimPol stimulation by PolDIP2. From both our data presented in the manuscript and the structural insight of PrimPol (Figure 2C), we propose the following model for PolDIP2-dependent PrimPol polymerase stimulation. The PolDIP2 region “A” in the T1 fragment interacts with the PrimPol regions “a” and “c”, in the vicinity of the dNTP binding pocket. PolDIP2 region “B” and “C” in the T2 and T3 fragment can interact with PrimPol regions “b2” and “b1”, respectively, which locate in the flexible loop of PrimPol (red). These multiple protein-protein interactions can be responsible for the function of the unique PolDIP2 Arg-cluster close to the pyrophosphate binding motif, (both present in T3 fragment), and also the T1 fragment of PolDIP2, likely contributing to optimize dNTP binding at the PrimPol 3’-elongation site, thus favoring processive DNA polymerization. This stimulation is likely crucial to facilitate elongation across DNA lesions.

In agreement with this idea, a more robust interaction with the primer/template could indirectly imply an increase in affinity for the correct incoming nucleotides. That would be very relevant, considering PrimPol’s inherent low affinity for dNTPs (38, 39) that could hinder its ability to synthesize DNA *in vivo*. Here we for the first time report a beneficial effect of PolDIP2 on PrimPol’s affinity for dNTPs (Figure 5A). Again, the improved dNTP binding depends on the T3 domain of PolDIP2. At this point, a relevant question is if the T3 domain of PolDIP2 can directly contribute to enhance dNTP binding by PrimPol. Detailed amino acid sequence alignments and 3D-modelling confirmed that the T3 domain of PolDIP2 shares a large similarity with bacterial ApaG proteins (Figure 3A and B). The function of ApaGs is currently unknown, but it has been proposed that they have the potential to interact with pyrophosphate moiety or even with complete nucleotides (26, 27). In addition to the pyrophosphate binding motif GxGxxG, the T3 domain of PolDIP2 has a cluster of proximal arginines that could be potential nucleotide binding ligands. In agreement with that, dismantling of the pyrophosphate binding motif, or alterations in the arginine cluster of PolDIP2 precluded the stimulation of dNTP binding (Figure 5C), and the consequent increase in polymerization by PrimPol (Figure 3 and 4).

In summary, we propose that the ability of the T3 domain of PolDIP2 to interact with the flexible loop of PrimPol triggers an improvement of dNTP binding which requires the ApaG-like pyrophosphate binding motif and its proximal Arg-cluster. Such a tighter binding is likely contributed by the T1 domain of PolDIP2, which is able to interact with regions “a” and “c” of PrimPol (Figure 6). Considering that region “a” is involved in the positioning of the templating base, whereas region “c” contains Lys297, a direct ligand of the pyrophosphate moiety of the 3’-incoming nucleotides (17), it is tempting to speculate that the T1 region contributes to the induced-fit step required to select the correct nucleotide.

Finally, we showed that at physiological relevant dNTP concentrations (< 12,5 *µ*M) PrimPol is a non-processive DNA polymerase, unable to incorporate more than a single nucleotide (Figure 5D), implying that PrimPol by itself is likely inactive *in vivo* as a polymerase. Association with PolDIP2 could activate PrimPol *in vivo* to overcome the relatively poor affinity for dNTPs and provide processivity. Moreover, PolDIP2 could boost the dNTP binding potential of PrimPol specially when the insertion has to occur opposite a templating base lesion. Future experiments with other PolDIP2-interacting DNA polymerases, such as Polλ and DNA Polη could show if this mechanism of stimulation by PolDIP2 can be extrapolated to other TLS DNA polymerases.

## Abbreviations

DNA: deoxyribonucleic acid
dNTP: deoxynucleoside triphosphate
nt: nucleotide
GST: glutathione S-transferase
IPTG: isopropyl β-D-1-thiogalactopyranoside
PBS: phosphate buffered saline
DTT: dithiothreitol
AEBSF: 4-(2-Aminoethyl) benzenesulfonylfluoride hydrochloride
ATP: adenosine triphosphate
EDTA: ethylenediaminetetraacetic acid
SDS: sodium dodecyl sulfate

## ACKNOWLEDGEMENTS

We would like to thank Mikael Lindberg from the Protein Expertise Platform from Umeå University for assistance with plasmids construction.

