## Supplementary Figure S1 for "A unique arginine cluster in PolDIP2 enhances nucleotide binding and DNA synthesis by PrimPol"

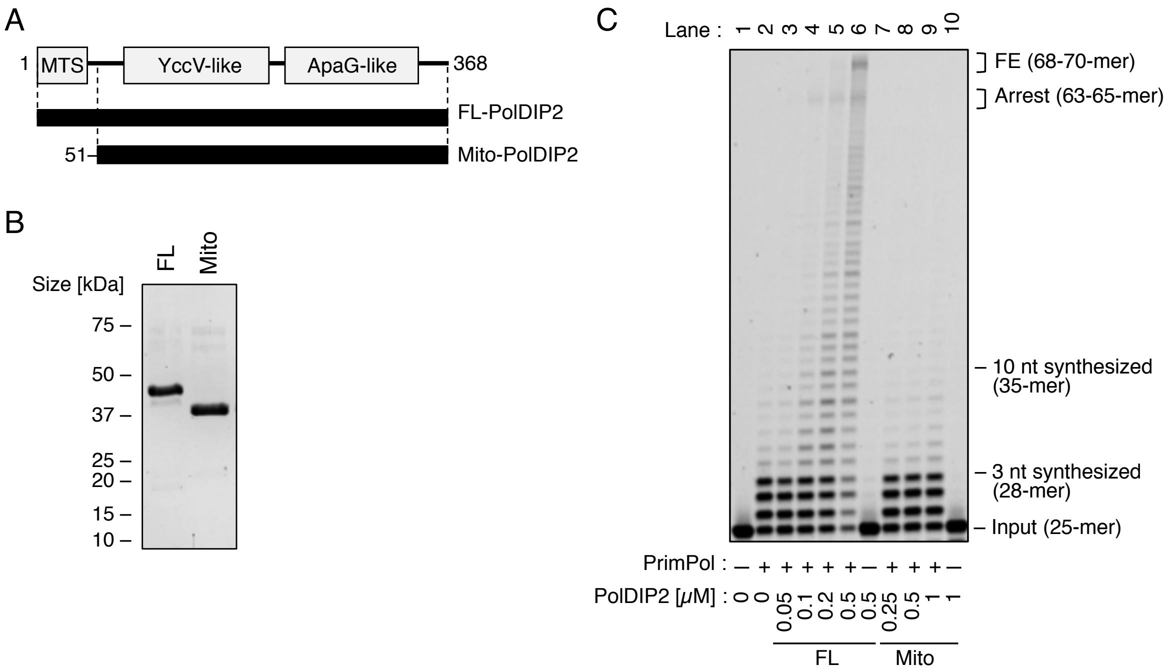


**Supplementary Figure S1.** Mitochondrial PolDIP2 is unable to stimulate PrimPol in DNA synthesis (A) Schematic presentation of the FL-PolDIP2 and the mitochondrial variant of PolDIP2 (Mito-PolDIP2). (B) Purification of Mito-PolDIP2. 300 ng of each purified PolDIP2 variant on SDS-PAGE gel stained with InstantBlue (Expedeon, UK). (C) PrimPol primer extension assays in the presence of FL- and Mito-PolDIP2. Reactions with 150 nM of PrimPol in the presence of 15 nM of 5'-TET-labeled primer/template DNA and the indicated amount of PolDIP2 variants.
