## Supplementary Figure S2 for "A unique arginine cluster in PolDIP2 enhances nucleotide binding and DNA synthesis by PrimPol"

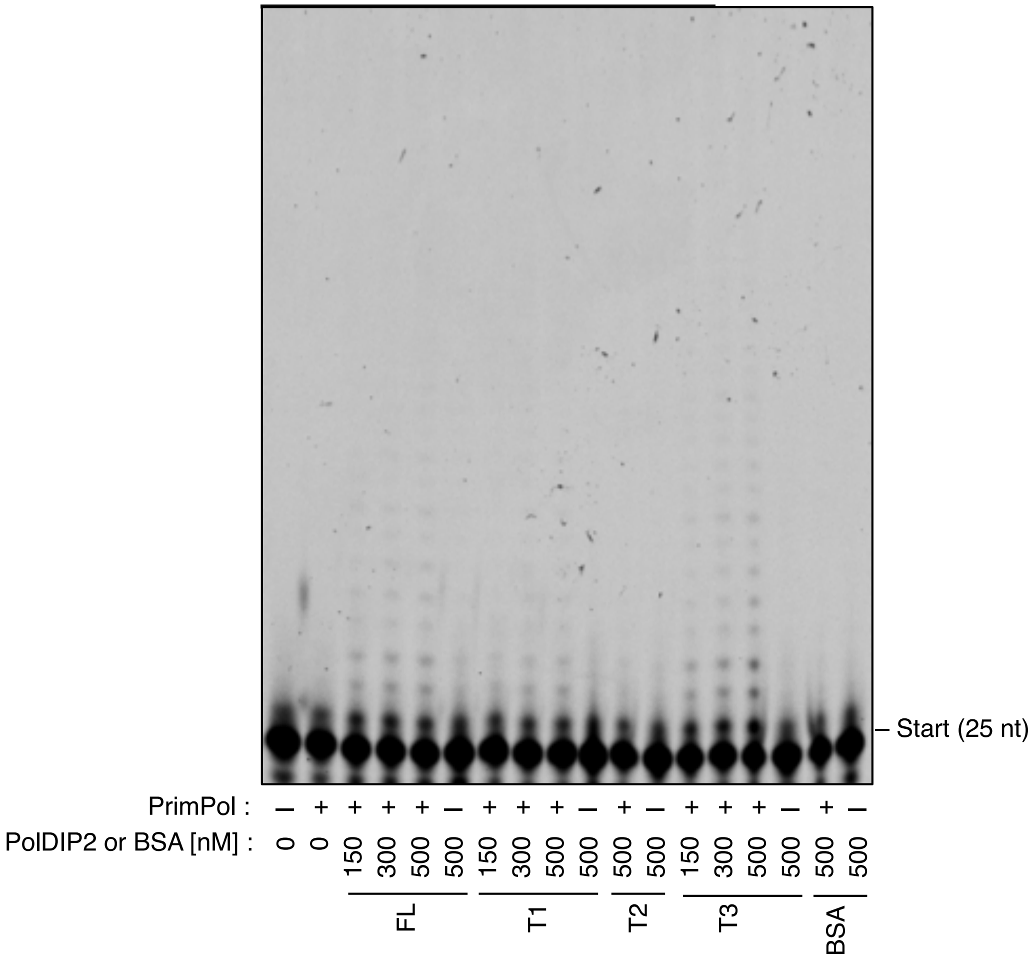


**Supplementary Figure S2.** PrimPol primer extension experiments in single hit conditions. Polymerization reactions in the presence of primer/template DNA (50 nM), a 10-fold excess molarity over PrimPol (5 nM) to minimize re-initiation of synthesis after a termination event.
