## Supplementary Figure S3 for "A unique arginine cluster in PolDIP2 enhances nucleotide binding and DNA synthesis by PrimPol"

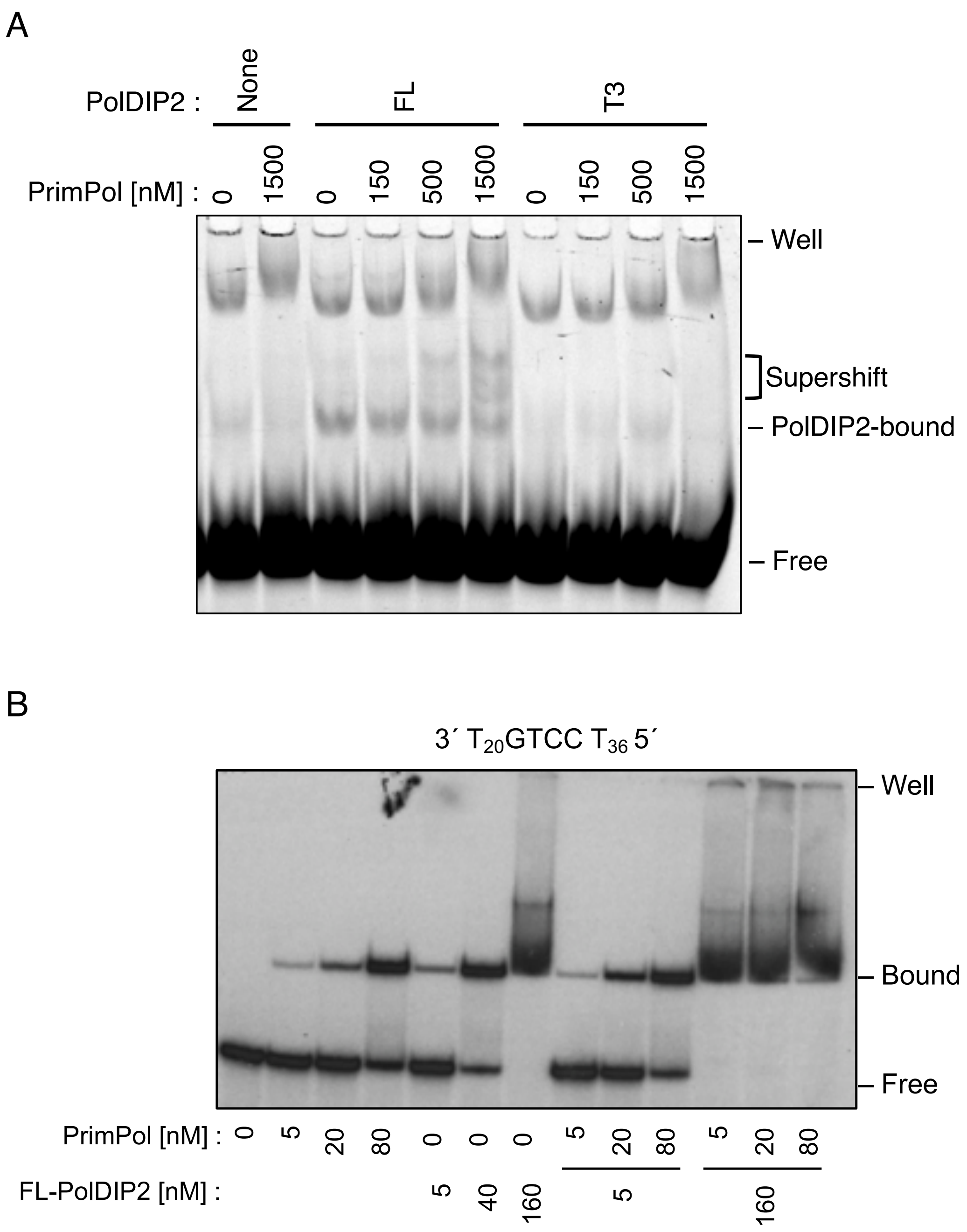


**Supplementary Figure S3.** Influence of PolDIP2 on PrimPol binding to different DNA templates, related to Figure 1H. (A) The T3 PolDIP2 fragment is unable to stimulate PrimPol-DNA binding. 15 nM of 5'-TET-primer/template DNA was mixed with the indicated concentrations of PrimPol and FL-PolDIP2 and incubated at 37 °C for 5 min. The PrimPol-dependent complexes are indicated with ‘supershift’. PolDIP2 bound to DNA (PolDIP2-bound), and unbound DNA (free) are indicated. (B) Interaction of PrimPol and PolDIP2 with ssDNA containing a favored PrimPol priming site (GTCC).

The binding affinity of PrimPol and PolDIP2 were compared by EMSA using the specific 60-mer GTCC template. 5 nM of $^{32}P$-labeled GTCC template was mixed with the indicated concentrations of PrimPol and FL-PolDIP2. PrimPol or PolDIP2 bound to DNA (PolDIP2-bound), and unbound DNA (free) are indicated.
