## Supplementary Figure S4 for "A unique arginine cluster in PolDIP2 enhances nucleotide binding and DNA synthesis by PrimPol"

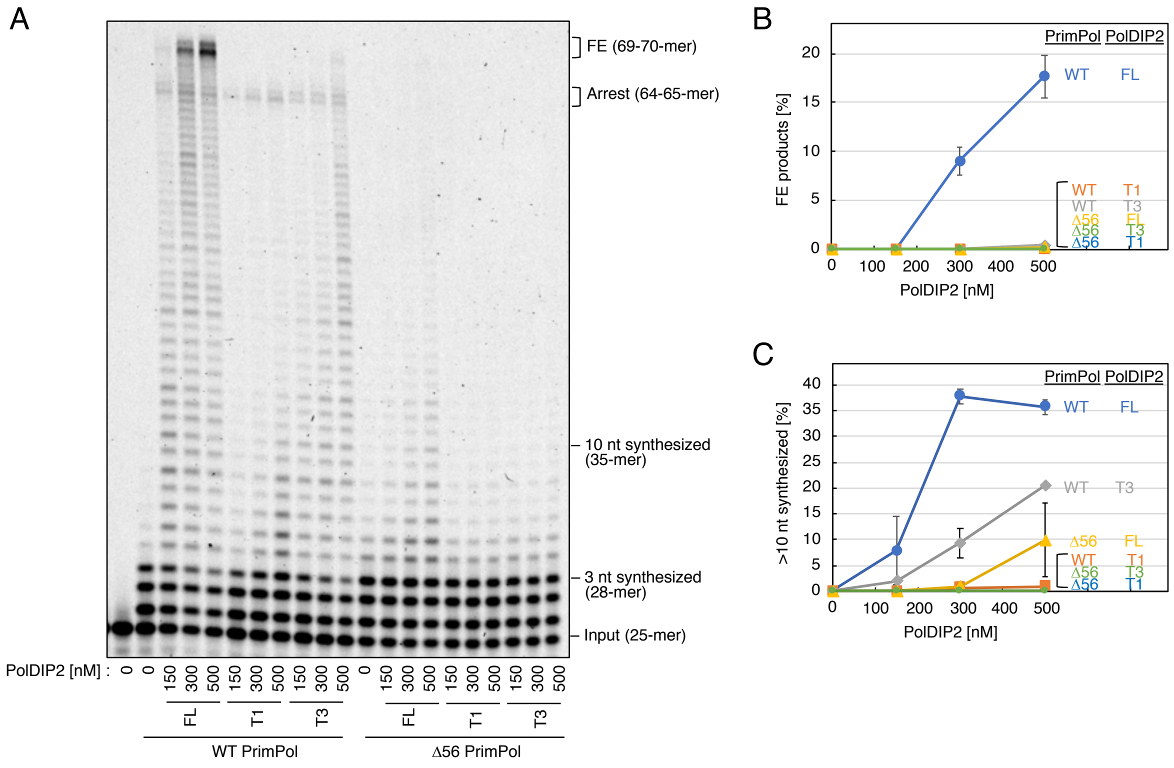


**Supplementary Figure S4.** AA 203-258 from PrimPol are required for PolDIP2-depended stimulation of DNA synthesis. (A) Primer extension reactions using 15 nM of 5'-TET-labeled template DNA and 150 nM of WT or Δ56 PrimPol incubated with the indicated amounts of PolDIP2 variants. Quantification of (B) full extension and (C) >10 nt synthesized DNA products. These experiments were performed twice, and averages with standard deviations are shown with error bars.
