## Supplementary Figure S5 for "A unique arginine cluster in PolDIP2 enhances nucleotide binding and DNA synthesis by PrimPol"

A

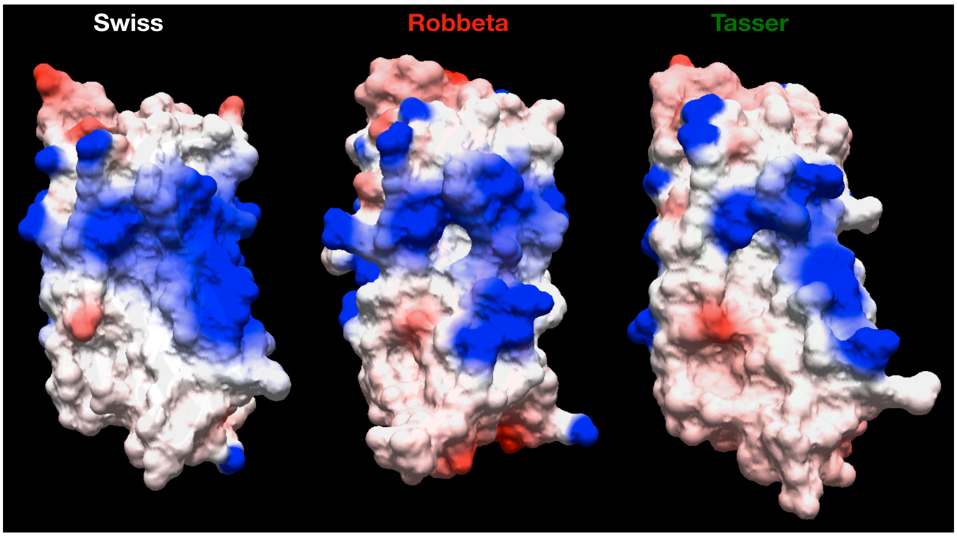


Z-score = –3.7 Z-score = –4.2 Z-score = –4.4

B

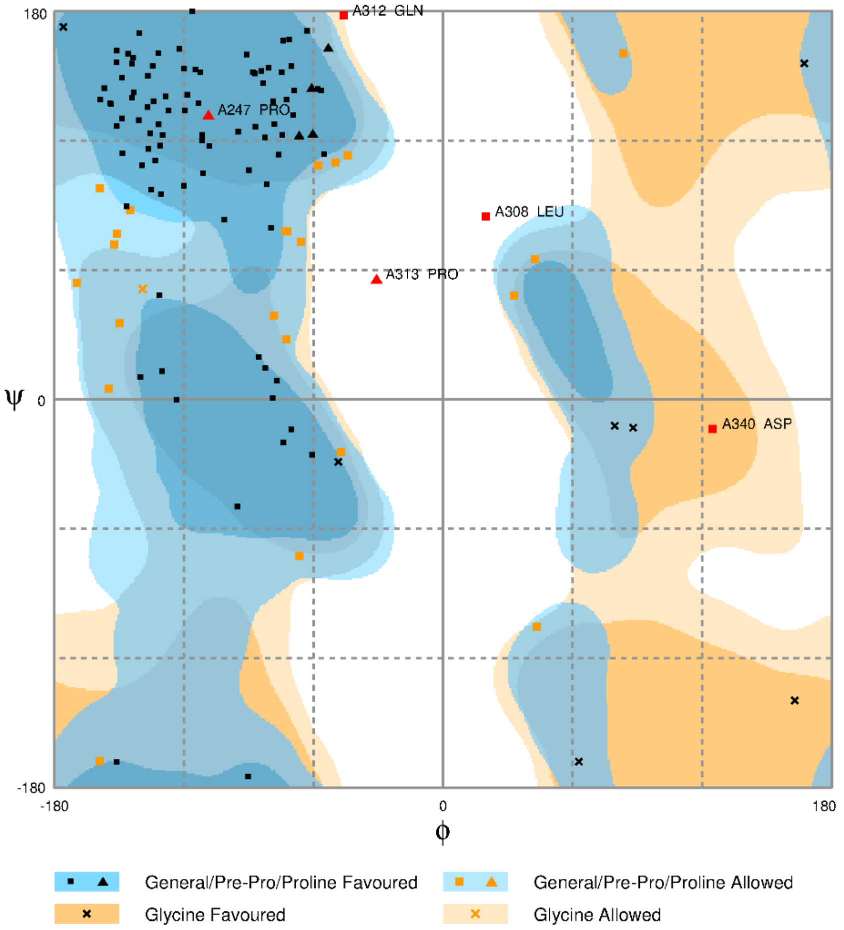


Number of residues in favoured region (~98.0% expected) : 100 (78.1%)
Number of residues in allowed region (~2.0% expected) : 23 (18.0%)
Number of residues in outlier region : 5 (3.9%)

**Supplementary Figure S5.** Structural model of PolDIP2-T3. (A) A Comparison of the *S. oneidensis* MR-1 ApaG (PDB id: 1TZA)-based modeled structures of PolDIP2-T3 with SWISS-MODEL, Robbeta, and I-TASSER web services, respectively. (B) The amino acid allowance of the constructed PolDIP2-T3 model (I-TASSER) was calculated with Ramachandra Plot.
