## Supplementary Figure S6 for "A unique arginine cluster in PolDIP2 enhances nucleotide binding and DNA synthesis by PrimPol"

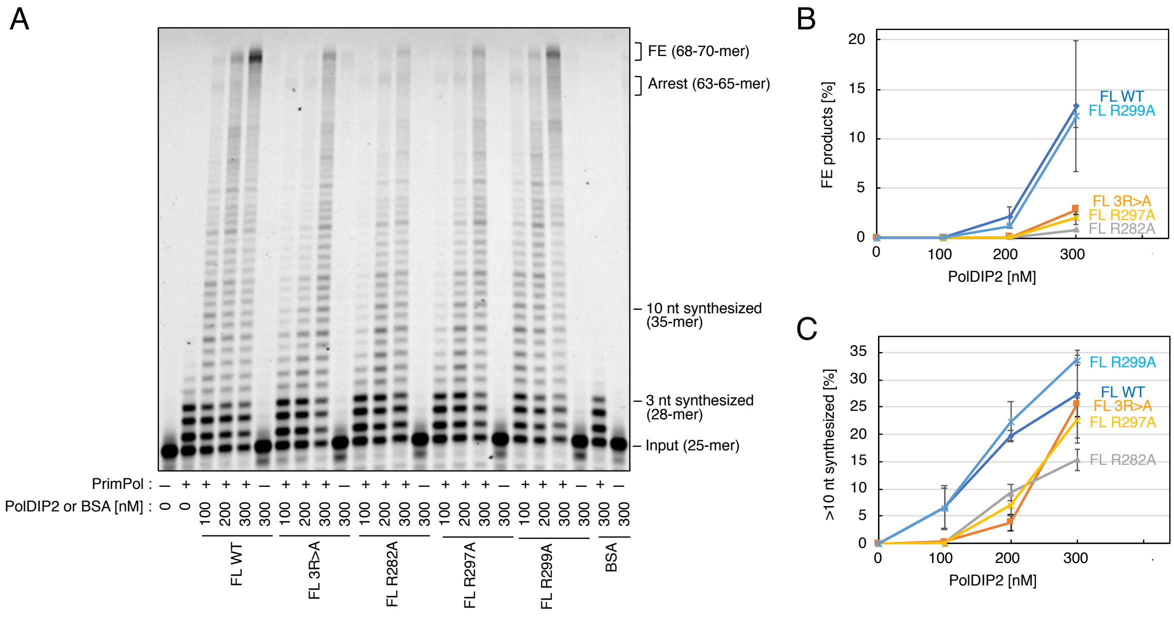


**Supplementary Figure S6.** The PolDIP2 Arg-cluster is required for PrimPol stimulation (A) Primer extension experiments using 15 nM of 5'-TET-labeled primer/template DNA and 150 nM of WT PrimPol in the presence of the indicated amounts of PolDIP2 variants. Quantification of (B) full extension and (C) >10 nt synthesized DNA products. These experiments were performed twice, and averages with standard deviations are shown with error bars.
